# Single exposure to social defeat or immobilization stress fails to induce lasting alterations in general anxiety and hippocampal neurogenesis in Wistar and wild-type Groningen rats

**DOI:** 10.1101/2020.02.12.946186

**Authors:** Deepika Patel, Ioannis Koutlas, Sebastiaan H. de Waard, Bauke Buwalda

## Abstract

Suppression of hippocampal neurogenesis is a readout for stress-induced alterations in neuroplasticity. In this study, we hypothesized that a single episode of severe social or non-social stress would differentially suppress neurogenesis in the dentate gyrus (DG) 10 days later in two rat strains. We anticipated that the suppression following social stress would be less severe in wildtype Groningen (WTG) rats, a rat strain considered relatively resilient to social stressors. Male Wistar and WTG were subjected to either social defeat or to immobilization stress. Behavioral response to social defeat and acute corticosterone response to both stressors was measured as well as anxiety behavior 10 days later on the elevated plus maze. Subsequently, brains were collected following cardiac aldehyde perfusion. The behavioral freezing response to defeat was much stronger in Wistar rats as compared to WTG rats. Acute corticosteroid response was similar in both strains although Wistar rats more rapidly resumed baseline values. There was no significant effect of both stressors on hippocampal DG cell proliferation and differentiation as well as on anxiety behavior. However, a striking strain difference appeared in anxiety behavior and both markers of neurogenesis. The WTG strain exhibiting much lower anxiety as well as reduced rate of hippocampal neurogenesis under all treatments. The results in this study suggest that both short-lasting acute stressors failed to induce lasting anxiety or decreased neurogenesis in the DG. Future studies could explore if and how rate of hippocampal neurogenesis is related with behavioral coping with stress.

## INTRODUCTION

Although exposure to severe stressful events potentially induces lasting changes in cognitive and emotional behavior major individual differences are reported in the susceptibility to stress-induced psychopathologies. For example, many people are experiencing a traumatizing event in their life but only a small proportion of these people develop a posttraumatic stress disorder (1) or a major depression (2). Previous experience and genetic makeup probably play an important role in these individual differences in susceptibility. Many preclinical studies study the neurobiology involved in the stress induced behavioral changes. Jacob et al. (3) formulated the neurogenic hypothesis of depression stating that the association between stress and depressive episodes is mediated by a decrease in neurogenesis in the dentate gyrus (DG) of the hippocampus. Whereas stress inhibits neurogenesis (4–6) in rats, the administration of SSRIs has been shown to increase it (4,7,8).

Although many studies focus on the detrimental consequences of repeated or chronic stress exposure there are clear indications that exposure to one or two short-lasting stressors can lastingly affect brain and behavior (9–11). Acute foot shock stress not only suppressed neurogenesis in the dentate gyrus immediately after the stress episode (12) but also up to one week later (13).

In a study by Mitra et al., (14) it was found that a single 2 hour lasting episode of acute immobilization stress was adequate to induce neuroplastic changes in the amygdala but only when this was measured 10 days after the episode and not when measured after one day. Here, we wanted to examine whether this neurobiological effect generalizes to brain regions other than the amygdala focusing on hippocampal neurogenesis. In our experiments we wanted to compare the effects of the non-social restraint stress with the ethologically relevant stress model of a single social defeat in the resident-intruder paradigm. We chose to use these two stress paradigms because both are frequently used in preclinical stress research to study mechanisms involved in the development of stress-related psychopathologies (15–17). Motta and Canteras (18) reported that the two stress models induced similar activation patterns in several brain regions, especially in those containing corticotropin-releasing hormone (CRH) neurons, but up to date there is no study comparing the effects of single exposure in the two models on neurogenesis. The exposure to stress of social defeat in general includes a physical interaction period of up to 15 minutes and a subsequent psychosocial stress period of 45 minutes without further physical attacks but threat of attack for the remaining time (19). Exposure to single restraint or immobility stress is usually applied for a period of 2 hours (14). Since we wanted to compare the lasting consequences of these two frequently used stress paradigms we decided to apply these different stressors, each with their own exposure time.

To quantify neurogenesis, doublecortin (DCX) immunohistochemistry was used. DCX is a microtubule-associated protein that is expressed in neuronal precursors making it a suitable marker of neurogenesis (20,21). As DCX marks young immature neurons, a specific maturation stage of the cell, it was used to gauge the survival of the new neurons (20). In order to distinguish between cell survival and proliferation, Ki-67, a marker of cell proliferation was used. Ki-67 is a protein that is expressed during the interphase and mitotic phase of the cell division cycle (22). To compare individual differences in susceptibility to these stressors we compared the results of the stress exposure in male rats of a typical laboratory Wistar rat strain with rats from a feral strain; descendants of rat pairs caught in the wild and outbred for many generations in our lab (23). Rats from the so-called wild-type Groningen (WTG) strain showed to be rather resilient to behavioral consequences of social defeat exposure in comparison with Wistar rats (24).

We hypothesized that stressed rats from both stress conditions would exhibit a significantly lower rate of neurogenesis compared to controls but that in comparison with WTG rats, Wistar rats would be particularly more prone to the effects of stress of social defeat.

## MATERIALS AND METHODS

### Animals and housing conditions

Male wildtype Groningen (WTG) rats, 8-9 weeks old, weighing 217 ± 23 g at the start of the experiment and male Wistar rats, 8-9 weeks old and weighing 305 ± 26 g were used and housed in groups of 5-6 animals before the start of experiment. Following the stress exposure, all animals were singly housed until they were sacrificed 10 days later (Fig. 1). Food and water were given *ad libitum*. All animals were housed under regulated lighting conditions (lights on at 10:00 h and off at 22:00 h). All experimental procedures were performed between 11:00h and 15:00h.

**Figure 1:**
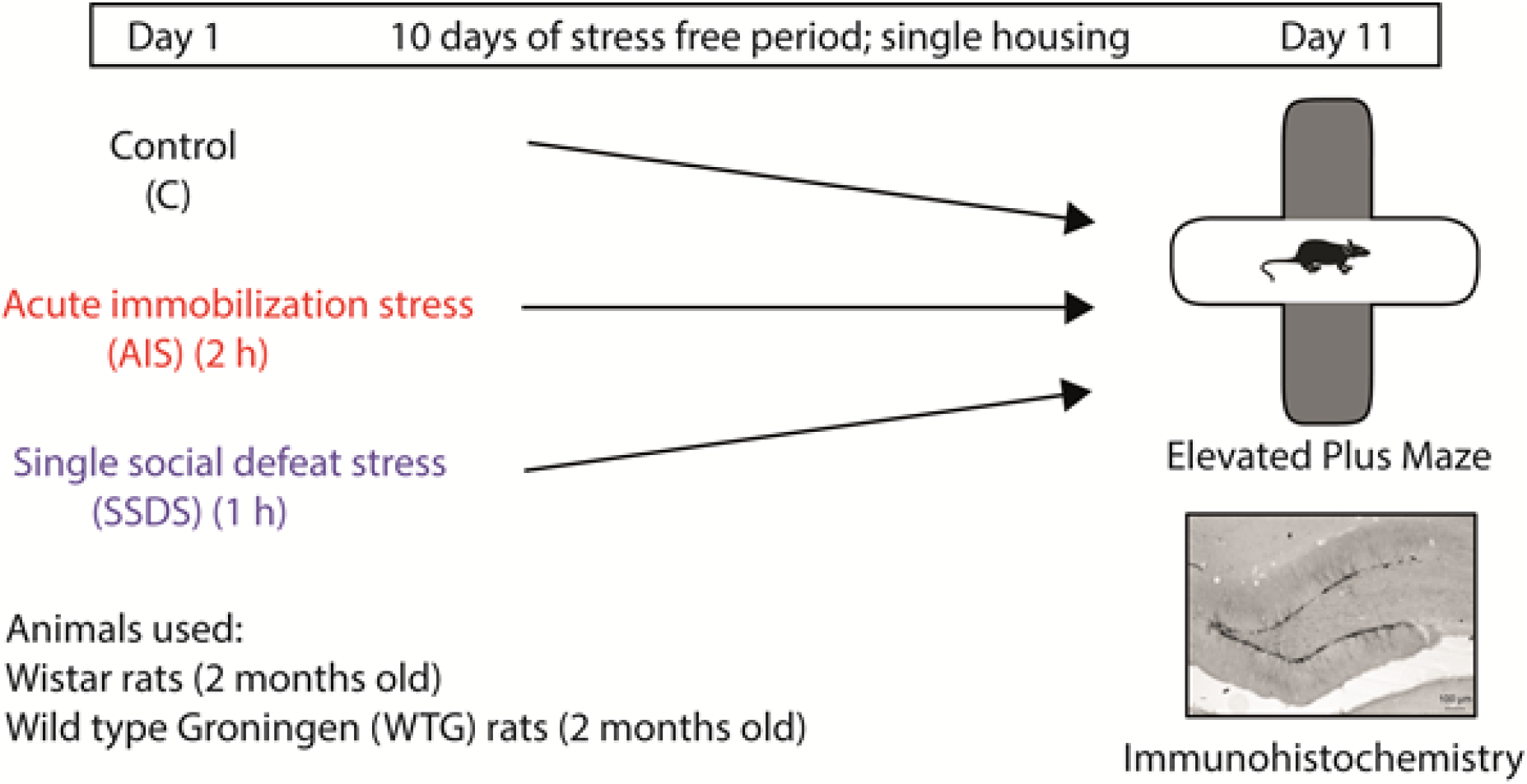
Timeline of the experimental procedure. Schematic representation of the experimental protocol in which rats (WTG and Wistar) were treated with acute immobilization stress (AIS), single social defeat stress (SSDS) or control procedure and tested for anxiety-like behavior after 10 days on elevated plus maze and 2 hours later the animals were sacrificed for further immunohistochemistry studies.

For the resident-intruder test the rats that were used as residents (i.e. dominant) were male WTG rats and approximately 6 months old. These rats were housed in observation cages (80 cm x 55 cm x 50 cm), together with an oviduct-ligated female to facilitate aggression and territorial behavior and to prevent social isolation. All experiments were approved by the Groningen University Committee on Animal Experiments (DEC 6746C).

### Stress paradigms

#### Immobilization

Wistars (N=8) and WTGs (N=8) were placed in a plastic bag that had a breathing hole at its point-shaped tip. The other end of the bag was tightened around the base of the rat’s tail. The animals were immobilized for a total of 2 hours (25,26). After two hours of immobilization, the animals were returned to their home cage and singly housed.

#### Resident-Intruder paradigm

The dominant WTG rats (N=23) were trained to attack intruder animals. After analyzing the training sessions, the most aggressive animals were chosen for the experimental trials. The cohabitating female of the dominant rat was removed one hour before the start of the experiment. Following that, Wistar (N=8) and WTG rats (N=8) were placed in the cage of the resident rat for 15 minutes after the initial attack (Fig. 2. A). Subsequently, the intruder was removed and put into a small wire mesh cage (30 cm x 14 cm x 14 cm), which was then placed back into the cage of the resident for another 45 minutes. This prevented that intruder rats were receiving further direct physical attacks but allowing sensory interaction with the residential male and psychosocial threat of attack (27). After having spent a total of 1 hour in the resident’s cage (15 min + 45 min), the intruder rats were removed and placed back in its home cage and singly housed.

**Figure 2:**
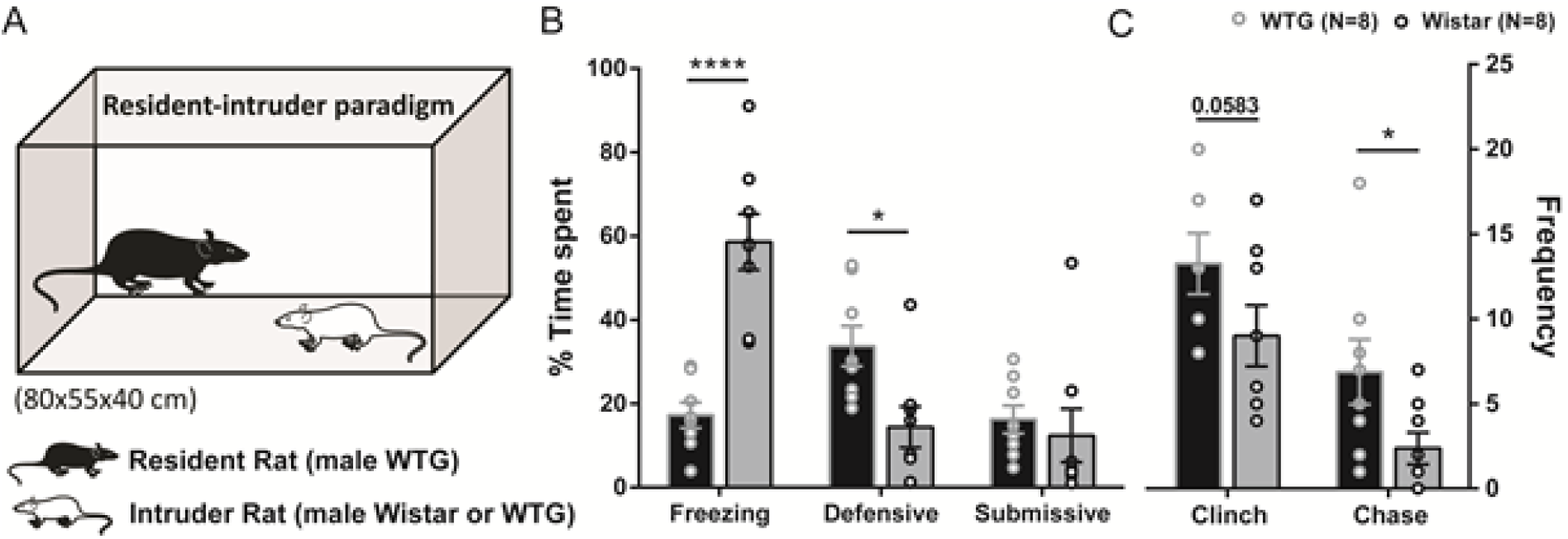
Behavior during the resident-intruder experiment. A. Schematic drawing of resident-intruder paradigm. B. Percentage of time spent on a number of selected behaviors during the resident-intruder experiment by the intruder rats (WTG and Wistar). C. Average number of times the intruder rats (WTG and Wistar) were attacked or chased. The black bars with grey circle indicates WTG rats and grey bars with black circles represents Wistar rats. Error bars expressed as mean ± SEM. Asterisks indicate significant differences (* p < 0.05, **** p < 0.0001 level, unpaired t-test).

### Behavioral analysis of resident-intruder experiment

The behavior of the socially defeated animals were recorded during the first 15 minutes of the confrontation. These videos were then analyzed. The displayed behaviors of the animals were scored using a homemade computer program (ELINE). For the different identified behaviors and their respective descriptions, see Table 1. The percentage of the time displayed by each behavior were calculated and are shown in figure 2 (B, C).

**Table 1.** Behaviors scored with the resident-intruder paradigm. The intruder rats from both the strains displayed several behaviors against the WTG residential male rat in the resident-intruder paradigm.

| Behavioral code | Behavior | Description |
| --- | --- | --- |
| Freezing | Freezing | No movement except for slight head-waving. |
| Follow/mirroring | Defensive | Following head of the resident to anticipate attack. |
| Upright |  | Defensive upright postures. |
| Submissive | Submissive | Supine position lying on the back. |
| Clinch | Clinch | Physical attack by the resident. |
| Chase | Chase | Running away from the resident. |

### Blood sampling

Before, during and after stress exposure blood samples were taken from the tip of the tail by tail clipping, to measure corticosterone (CORT) levels in the blood. This was performed on all treatment groups. Collection of each blood sample took on average about 30 sec −1 min. A total of 5 samples per animal were taken that were cooled in ice water. A baseline blood sample were taken prior to each stress paradigm (t = −10min). A second blood sample was taken right after the physical interaction period with the resident prior to placing socially stressed rats into the wire mesh cage (t = 15min) or after 15 minutes of immobilization in the plastic bag. The third blood sample was taken after completion of resident intruder paradigm (for socially stressed rats) and at the middle of the immobilization stress (rats subjected to AIS) (t = 60min). At t = 120min, a blood sample was collected at the end of the immobilization experiment and 1 hour after end of the social stress period (reflecting recovery in the social defeat experiment). Finally, blood was sampled to measure recovery following immobilization stress (t = 180min). Ten μl heparin was added to each sample to prevent clotting. The samples were then centrifuged and plasma was collected and stored at −20 °C until the corticosterone assay was performed.

### Analysis of plasma corticosterone

Corticosterone levels were determined by radioimmunoassay (MP Biomedicals, Orangeburg, NY) with an intra-assay variation of 7.2% and an inter-assay variation of 6.9%. Samples were analyzed *in-duplo* and measures were averaged afterwards.

### Elevated plus maze test

Ten days after the experiments all treatment groups were tested for anxiety-like behavior on an elevated plus maze (height: 50 cm from the ground; length of arms: 45 cm). The test was performed between 10 AM and 12 PM, and each animal was tested for 5 minutes. Light intensity on an open arm was around 80 lux, and on the closed arm between 2 and 5 lux. Each animal was taken separately to the room with the elevated plus maze and placed in the middle of the maze, facing towards an open arm (Fig. 4. A). Individual trials were videotaped for subsequent analysis. In between two trials, the maze was cleaned with warm water and soap and dried afterwards. The relative amount of time spent on an arm was quantified using the behavioral analysis program ELINE. Time spent on a specific arm was only scored when the animal was on that arm with all four paws. The performance was assessed by calculating the relative percentage of the total time each animal spent on the open arms [time on the open arm/(time on the open arms + time on the closed arms)].

**Figure 3:**
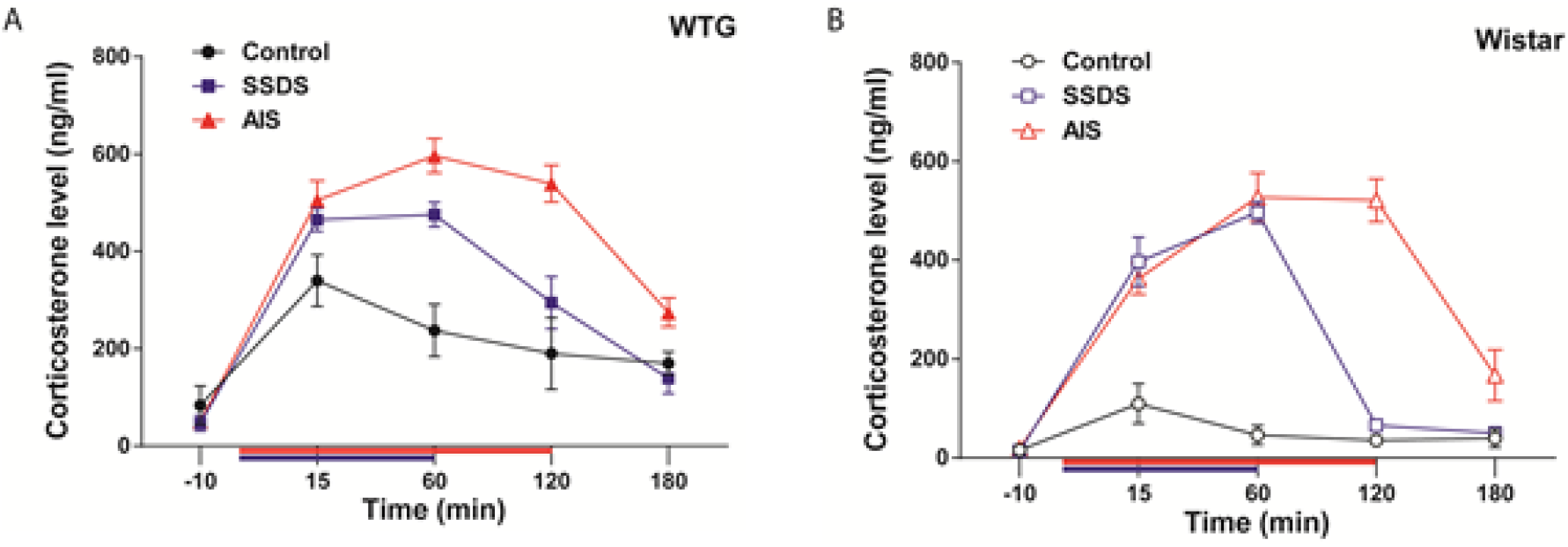
Plasma corticosterone responses in the rats. A. illustrates responses in WTG rats and B. represents data from Wistar rats with different treatments (control, SSDS, AIS). The average corticosterone levels per treatment (ng/ml) were measured at 5 time points (−10, 15, 60, 120, 180 min). Duration of actual stress exposure is indicated on the X-axis (red: AIS and blue: SSDS).

**Figure 4:**
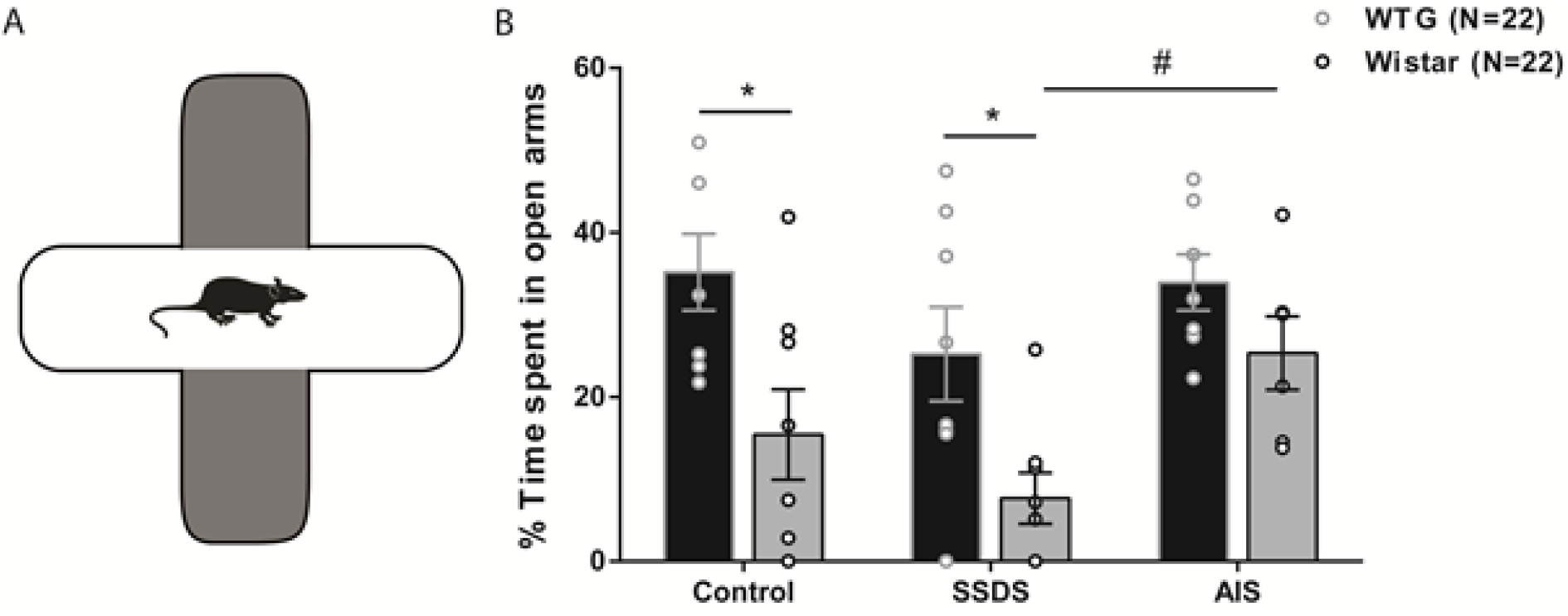
A. Schematic image of the elevated plus maze to study anxiety-like behavior after the animals being subjected to a single episode of acute stress (AIS and SSDS). B. Comparison between the performance of both strains (WTG and Wistar) on the elevated plus maze measured in % time spent on open arms [=time on open arms/ (time open arms + time closed arms) x 100]. Error bars expressed as mean ± SEM. Asterisk and hashtag indicates significant differences (* p < 0.05, # p < 0.05 level, Sidak’s test for multiple comparison).

### Sacrificing and preparation of tissue

Animals were anesthetized with an overdose of intraperitoneally injected sodium pentobarbital and subsequently perfused using a solution of heparinized saline (10 ml heparin/L saline) for approximately 1 minute followed by a 4% paraformaldehyde solution in 0.1M phosphate-buffer saline (PBS), 250 ml per animal. The brains of the animals were removed and placed in 0.1 M PBS solution, then placed in a 30% sucrose solution for 24 h to prevent ice crystals forming in the tissue. Finally, the brains were frozen and stored at −80 °C until coronal sectioning (30 μm) with a cryostat. From each animal similar dorsal hippocampus sections (bregma −3.00 to −3.96 according to paxinos & Watson, 2007) were selected for immunohistochemical staining of DCX and Ki-67. Immunopositive cells were counted using an Olympus BH-2 optical microscope at 40x magnification.

### Immunohistochemistry

#### Doublecortin (DCX) staining

The number of young differentiating neurons were identified with an antibody against DCX (1:1000; polyclonal goat-anti-DCX; Santa Cruz Biotechnology, Santa Cruz, CA, USA). The sections were incubated with the primary antibody in 0.01M PBS for 3 days at 4°C. Following that sections were incubated with biotinylated rabbit-anti-goat immunoglobulin (1:500, Jackson ImmunoResearch Laboratories, Inc., West Grove, PA, USA) overnight at 4°C. Amplification was achieved with avidin-biotin complex (1:500, ABC; Vector Laboratories, Peterborough, UK) for 2h at room temperature. Finally, chromogen development was performed with diaminobenzidine (DAB, DAB substrate kit, Roche Diagnostics) and all the sections were mounted on glass slides, dehydrated with a graded ethanol-xylol treatment and embedded in DPX-mounting media.

For quantification of the DCX immunopositive cells, the area around the infrapyramidal blade of the dentate gyrus was selected bilaterally and the proportion of area covered with stained cell bodies to the total selected area constituted our final variable. Moreover, the stained cell dendrites were quantified in a similar way by selecting the area of the molecular layer ventral to the infrapyramidal blade. The dendrites and cell bodies were quantified separately because different intensity thresholds had to be used. The DCX sections were analyzed using a Leica DMI 6000B optical microscope at 40x magnification. Quantification was performed using LAS Image Analysis Software (Leica, Cambridge, UK).

#### Ki-67 staining

For the detection of proliferating cells, an antibody against Ki-67 protein was used (1:200; mouse-anti-Ki-67; Monosan; Uden, The Netherlands). The incubation time for the primary antibody was 40h in 0.01M PBS at 4°C. Next, the tissue were incubated with biotinylated goat-anti-mouse immunoglobulin (1:400; Jackson ImmunoResearch Laboratories, Inc., West Grove, PA, USA). Amplification was achieved with avidin-biotin complex (1:400, ABC; Vector Laboratories, Peterborough, UK) for 2h at room temperature. Chromogen development was performed with diaminobenzidine (DAB, DAB substrate kit, Roche Diagnostics) and all the sections were mounted on glass slides using a gelatin solution. A treatment with Mayer’s hematoxylin solution was used to counterstain the tissue. Finally, the tissue were dehydrated with a graded ethanol-xylol treatment and embedded in DPX-mounting media.

### Statistics and data presentation

Data provided are expressed as group mean ± standard error of measurement (SEM). Statistical analysis was performed using GraphPad Prism 6. Two-way analysis of variance (ANOVA) was performed to compare immunohistochemistry data across treatments (social defeat, immobilization and control), strain (WTG/Wistar), and the interaction of these factors followed by *post hoc* Sidak’s test for multiple comparison. Unpaired t-test was used to compare the behavior scored during the resident-intruder paradigm. For the corticosterone measures repeated measures was used followed by Tukey’s multiple comparisons test. The significance level was set to *p* < 0.05.

## RESULTS

### Behaviors scored during the resident-intruder experiment

Behaviors displayed by the socially defeated rats (WTG and Wistar) were scored and pooled into 3 categories (as described in table 1). The data is presented in the figure 2. B and C. The results show a striking difference in the behavioral response to the aggressive behavior of the residential male between WTG and Wistar rats. Freezing behavior in Wistar rats (58.61 ± 6.657, N=8, *p* < 0.0001) is more than 3 times higher than in WTG rats (17.25 ± 3.030, N=8). In contrast, WTG rats (33.71 ± 4.800, *p* = 0.0137) depicted significantly more defensive behavior compared to defeated Wistar rats (14.38 ± 4.905). No significant difference was observed between the strains considering submissive behavior.

The total frequency of attack and chase incidents were also scored and analyzed. Unpaired t-test pointed out that WTG rats (13.25 ± 1.800, *p* = 0.0583; 6.875 ± 1.913, *p* = 0.0267 respectively) were attacked and chased significantly more than the Wistar rats (9.000 ± 1.793, 2.375 ± 0.9437 respectively).

### Neuroendocrine stress response: differences in plasma corticosterone response between treatments and strains

Considering WTG rats, the differences in corticosterone response per treatment becomes apparent after the first 15 minutes of the experiment. From t=60 min, the corticosterone response during immobility treatment is higher than after the defeat exposure. WTG rats also show a significant response to the control procedure. Analysis by two-way repeated measure (RM) ANOVA with treatment and time as variables indicate a significant interaction between these two variables (F (8, 64) = 6.900, *p* < 0.0001). *Post-hoc* analysis with Tukey’s multiple comparisons test showed significant increase in CORT levels in AIS treated rats compared to the control rats (at 15 min). At t = 60 min (when the social stress exposure ends) corticosterone levels in all three treatments are significantly different from each other (control vs. SSDS: *p* = 0.0002; control vs. AIS: *p* < 0.0001; SSDS vs. AIS: *p* = 0.0348). At t = 120, significant difference was seen between control vs. AIS *(p* < 0.0001) and SSDS vs. AIS *(p* < 0.0001). At t = 180 min, indicating recovery for all procedures, only SSDS vs. AIS shows significant difference with *p* = 0.0156. Overall, it can be concluded that the difference of the immobilization treatment in the WTG in comparison with the defeat and control treatments is strikingly high. The overall response of the socially defeated group was also higher than that of the control WTG rats (Fig. 3. A).

Regarding the Wistar rats, similar effects are seen although some striking differences appear in comparison with the response of the WTG rats. Till t = 60 min corticosterone response to immobility and defeat are similar. Repeated measure ANOVA shows a significant interaction between treatment and time (F (8, 60) = 23.39, *p* < 0.0001) (Fig. 3. B.). The hormonal response to the control procedure is much lower than observed in the WTG rats (significant at t = 15 and 60 min *post-hoc* Tukey’s multiple comparison test *p* < 0.05). Speed of recovery after defeat stress (decline in corticosterone levels from t = 60 min to t = 120 min) and immobility stress (decline in corticosterone levels from t = 120 min to t = 180 min) is, however, different between strains. WTG rats show a more gradual downward slope of the corticosterone curve (a slower recovery to baseline) with both stressors (SSDS *(p* = 0.0004) and immobilization *(p* = 0.0025)) reflecting a reduced speed of recovery.

### No lasting effects of stress on anxiety like behavior 10 days after stress exposure but Wistars are more anxious than WTG rats

All the animals were behaviorally tested on the elevated plus maze test prior to brain collection (Fig. 4.). No lasting effects on anxiety behavior were observed in Wistar or WTG rats 10 days after either immobility stress or social defeat stress in comparison with the controls. Comparing both strains showed a clear difference in anxiety like behavior. WTG rats overall were less anxious on elevated plus maze compared to the Wistar rats, with significant differences for the control and social defeat groups (*p* = 0.0133; *p* = 0.0248 respectively). When it comes to comparison of the immobilized groups, significant strain differences disappeared.

### Hippocampal cell proliferation and neurogenesis is higher in Wistar compared to WTG rats

#### Hippocampal neurogenesis

To study the impact of different stressors on adult neurogenesis, young granule cells were stained with an antibody against doublecortin (Fig. 5). Two-way ANOVA revealed no significant effect of stress condition on DCX immunolabeling (F(2, 39) = 0.3323, *p* = 0.7193). The percentage of the area of interest covered by DCX-immunopositive cell bodies did not differ significantly between control, social defeat and immobilization conditions. Similar results were obtained when examining the effects of condition on the percentage of the area of interest that was covered with DCX-immunopositive dendrites (plot not shown). In essence, a single stress episode (social defeat or immobilization) did not have an effect on the number of cells expressing DCX.

**Figure 5.**
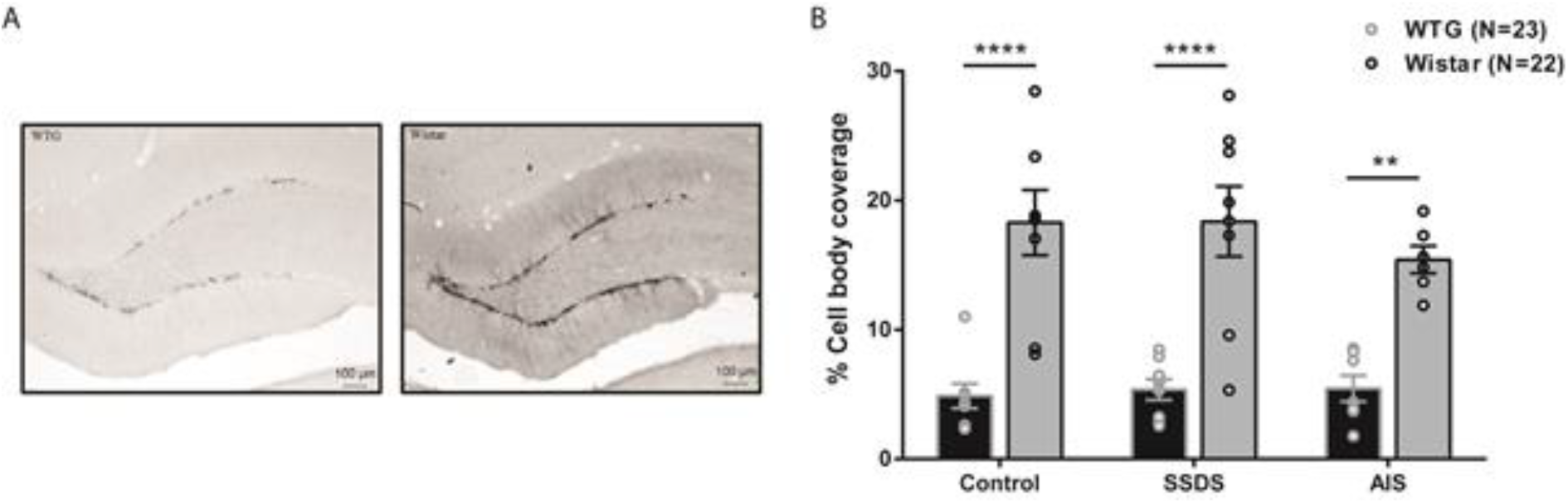
Effect of stress on cell survival in the dentate gyrus (DG). A. Photomicrographs of the DG of both strains (WTG and Wistar) control group, stained for DCX are shown. B. Differences in the rate of neurogenesis between Wistar and WTG rats. The black bars with grey circle indicates WTG rats and grey bars with black circles represents Wistar rats. Error bars expressed as mean ± SEM. Asterisks indicate significant differences between the strains (** p < 0.01, **** p < 0.0001 level, Sidak’s test for multiple comparison).

Comparing strains in each conditions, the test identified a significant effect of strain regarding cell bodies (F(1, 39) = 68.46, *p* < 0.0001) and dendrites. More specifically, Wistar rats had a significantly higher percentage of area covered with DCX-immunopositive cell bodies compared to WTG rats (control: *p* < 0.0001; SSDS: *p* < 0.0001; AIS: *p* = 0.0022).

#### Hippocampal cell proliferation

To investigate effect of two different stressors and strains on cell proliferation, quantification of Ki-67 positive cells in the dentate gyrus was performed. As shown in figure 6 no significant differences between treatments were found in the number of Ki-67+ cells when tested with two-way ANOVA (F(2, 34) = 2.283, p = 0.1174). The average number of Ki-67 positive cells in the DG did not differ significantly between control, social defeat and immobilization conditions. It appears that a single episode of stress (social defeat, immobilization) failed to produce a significant effect on the number of proliferating cells in the DG.

**Figure 6.**
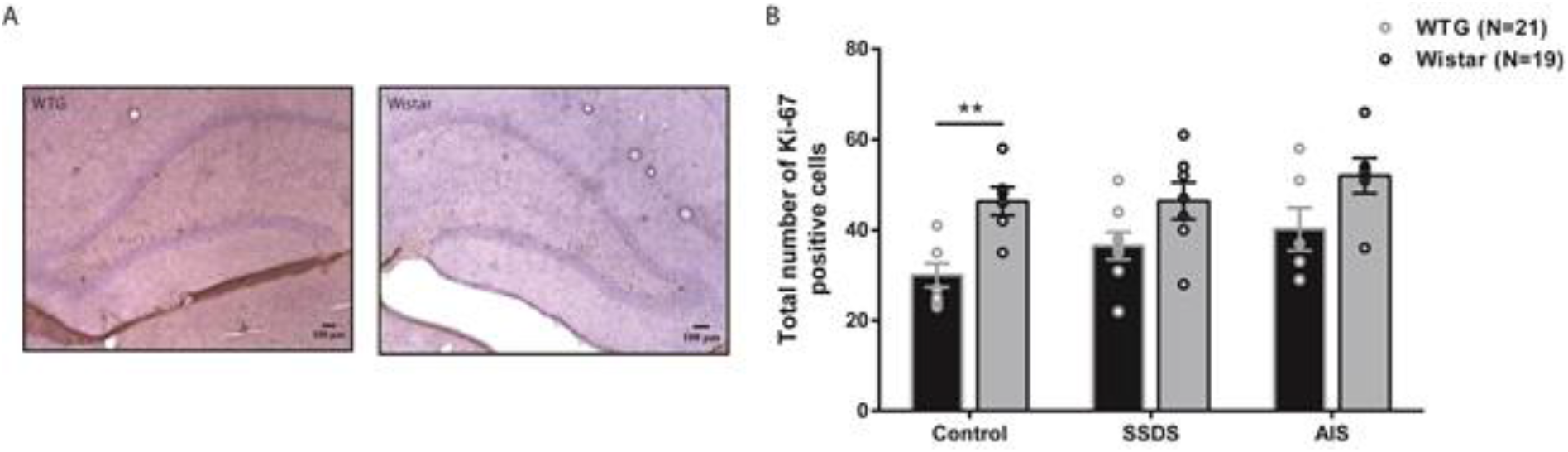
Effect of stress on hippocampal cell proliferation. A. Photomicrographs of the DG of both strains (WTG and Wistar) control group, stained for Ki-67 (marker for proliferating cells). Positive cells can be seen along the granular cell layers of the supra- and infra-pyramidal blade of the dentate gyrus. B. Difference in the cell proliferation between Wistar and WTG rats. The black bars with grey circle indicates WTG rats and grey bars with black circles represents Wistar rats. Error bars expressed as mean ± SEM. Asterisks indicate significant difference between the strains in the control group (** p < 0.01, level, Sidak’s test for multiple comparison).

A significant effect of strain on the number of Ki-67 positive cells was, however, identified (F(1, 34) = 18.54, p = 0.0001) with post-hoc testing indicating that control Wistar rats (p = 0.0097) had a significantly higher number of Ki-67 cells compared to WTG rats.

## DISCUSSION

In this study, we have investigated the efficacy of acute exposure to an aversive social or non-social experience in modulating the behavioral, physiological and neuroplastic response in two different rat strains, WTG and Wistar rats. To this end, we examined the effect of two frequently used animal models to induce stress being either a single episode of immobilization (2 hours) or social defeat (1 hour) on the acute corticosterone response and on anxiety behavior, neurogenesis and cell proliferation 10 days later.

The behavioral response of Wistar and WTG rats to the aggressive residential male was very different with Wistars showing mainly freezing behavior and WTG rats showing a lot more defensive-like behaviors. The corticosterone response to treatment was also different in WTG and Wistar rats with WTG rats showing a relatively strong response to the mild stress of the control procedure (placement of the home cage in an unfamiliar empty room). The peak levels reached after AIS and SSDS exposure probably reflect ceiling effects. This ceiling is reached in WTG rats exposed to immobility stress but not in social defeat stress whereas the peak levels to AIS and defeat were reached and therefore similar in Wistar rats. This may indicate that WTG rats are less affected by the exposure to the social aggressive behavior of the residential male than Wistar rats. This probably is related to the stronger defensive behavioral response in WTG rats. Previously it was shown that rats that resist to defeat by showing defensive behaviors are less vulnerable to the lasting consequences of social defeat stress (28). It seems that WTGs exhibit a lot more ‘proactive behaviors’, while Wistars exhibit a lot more ‘reactive behaviors’ (29,30). Even though the WTG rats seem to cope better with the social defeat experience, the speed of recovery of the neuroendocrine stress response is delayed in comparison with that in Wistar rats. This delayed recovery is also observed in WTG rats after AIS. Exposure to AIS offers no real coping strategy and therefore the neuroendocrine stress response is probably similar in both strains. The slower recovery in WTG rats is somewhat unexpected since WTG rats are reported to be more resilient to at least the social stress exposure (24). We postulated previously that the initial height of the response curve doesn’t indicate the degree of uncontrollability, instead the downward recovery slope does (31). In that respect it is surprising to see the reduced recovery slope in WTG rats. It is possible, however, that this relation between speed of recovery and the perception of controllability does not hold for a single exposure to a stimulus. In a comparison of the physiological and corticosteroid response during and after exposure to social defeat and a mating opportunity with a sexually receptive female it appeared that speed of recovery was faster after a single exposure to defeat in male rats than in males allowed to copulate once with a female. Only after repeated exposures, the recovery in the non-aversive social stimulus, sex, became faster than that following defeat (32). Not only the difference in speed of recovery of the corticosterone response is striking, also the response to the relatively mild stress of the control procedure is very different. WTG rats are much more sensitive with regards to the endocrine response to the transfer of the home cage to an unfamiliar room. Indeed, arousal in WTG rats is much more easily triggered in our experience than in Wistar rats. One could hypothesize that frequent arousal induced corticosterone secretion during the housing period prior to the experiment plays a role in the reduced hippocampal neurogenesis in WTG rats but this is difficult to confirm in this study. In the current experiments rats of both strains were singly housed and further left undisturbed after the stress exposure.

Even though robust corticosterone responses were measured during and after the acute stressors in the animals, we found no significant effect of stress on anxiety behavior as measured 10 days later on the elevated plus maze and in neurogenesis as measured with a cell proliferation (Ki-67) and a cell survival (DCX) marker. However, a significant strain difference was found with Wistar rats showing more anxiety-like behavior along with increased expression of DCX and Ki-67 as compared to the WTG rats, the strain regarded as being relatively resilient to social stress. Since anxiety is reported to correlate negatively with hippocampal cell survival and proliferation (33) the strain differences in anxiety and hippocampal neurogenesis are unexpected.

Although numerous studies suggest that inducing stress is able to suppress neurogenesis, the findings are nonetheless inconclusive. More specifically, while inducing multiple episodes of stress has been in most cases successful in suppressing neurogenesis (4,5,34–36), inducing acute stress has produced mixed findings. For example, while some studies have found that acute stress is able to suppress cell proliferation after time intervals similar to the one used in this study (12,13), a few studies have suggested that several acute stress paradigms are unable to induce neuroplastic changes in the DG (37,38). More specifically, when Dagytė and colleagues induced stress to rats using a single episode of restraint stress or the resident-intruder paradigm they found no change in cell proliferation in the DG after one, seven or 21 days. Contrastingly, Kirby and colleagues (39) found that inducing acute stress by means of immobilization was associated with increased cell proliferation in the dentate gyrus 3 hours after the stress episode and that this finding correlated with increased levels of corticosterone.

It was previously reported that a single episode of 2 h of immobilization stress caused up-regulation in spine density in the basolateral amygdala 10 d later along with significant increase in anxiety like behavior (14). We were unable to confirm the increase in anxiety in our experiments. Wistar rats subjected to immobilization stress were even significantly less anxious than the socially defeated Wistar rats. Among the WTG rats, the amount of anxiety on the elevated plus maze of the stressed animals is the same as that of the controls.

The main finding from this study challenges the hypothesis that acute stress can lastingly suppress neurogenesis through the action of glucocorticoids (40). Our study showed that the number of newly formed cells 10 days after acute stress is not associated with elevated corticosterone values or anxiety as measured by the EPM. It seems, that peak levels in plasma corticosterone levels after acute stress exposure, is not necessarily sufficient to suppress neurogenesis. Several factors might play a role in this such as the type of stress applied, time interval and procedures followed between stress and sacrifice, as well as strains used for the study including individual differences between the animals.

## ACKNOWLEDGEMENTS

This work was supported by the Groningen Institute for Evolutionary Life Sciences (GELIFES), Groningen, The Netherlands. We thank Jan Keijser for his assistance in imaging and image analysis.

